# Neurotropic bornaviral pressure as a potential driver of mammalian encephalization

**DOI:** 10.64898/2025.12.02.691862

**Authors:** Emmanuel Gauvain

## Abstract

**Summary:** Mammalian encephalization has increased episodically and unevenly across lineages, and classical ecological or social explanations account for only part of this variation. Because stable social groups facilitate the circulation of persistent neurotropic viruses, we asked whether long-term viral exposure may have contributed to shifts in brain-body scaling. Using published records of endogenous bornavirus-like elements (EBLs, genomic remnants of ancient neurotropic RNA viruses), we show that mammalian clades with steeper encephalization slopes tend to exhibit higher EBL burdens, whereas highly social lineages without pronounced brain expansion show minimal evidence of past bornaviral integration. These patterns suggest that viral exposure may modulate the association between sociality and encephalization.

To explore potential mechanisms, we modeled the evolutionary dynamics of *ARHGAP11B*, a hominin-specific gene that promotes basal progenitor amplification. Under weak cognitive selection alone, fixation of such an allele is slow and unlikely; however, under episodic, bornavirus-like developmental stress, fixation becomes substantially more probable when the allele is present as standing variation. This suggests that persistent neurotropic viral pressure could accelerate selection on neurogenic pathways.

Together, these findings motivate a broader consideration of neurotropic viruses as intermittent selective agents that may interact with social structure and development to shape long-term patterns of mammalian brain evolution.

## Introduction

Mammalian encephalization (i.e., increase in relative brain size with respect to body mass) has arisen repeatedly but unevenly across evolution, despite the important energetic cost of building and maintaining nervous tissue ^1^. Ecological and behavioral frameworks such as the cognitive-buffer ^2^, socioecological ^3^, and social-brain hypotheses ^4^ help explain broad interspecific variation, but struggle to account for the bursty, lineage-specific nature of mammalian brain evolution, thus impeding our understanding of the significant selective forces behind the evolution of complex nervous systems.

Notable work by Shultz and Dunbar (2010) reframed the problem around temporal changes in relative brain size (“encephalization slope”) within taxa, showing that encephalization trends are tightly associated with long-term social stability and cohesive group-living ^5^. However, some outlier taxa outside the scope of their research, especially among rodents and bats, exhibit varying and puzzling encephalization trajectories relative to social structure ^6^. Moreover, so far, no well-supported environmental factor has been linked to the selective pulses necessarily added to longer-term low-level pressures ^7^. More broadly, too little is known about the mechanisms through which such social environments generate strong, recurrent selection on neural development.

One important but often overlooked aspect of stable social groups is increased persistent viral exposure, especially to pathogens that chronically infect the developing nervous system. Neurotropic RNA viruses circulate efficiently in group-living mammals, yet the potential evolutionary impact of such infections on brain development has received little attention. Among them, bornaviruses (BDV) stand out as uniquely tractable: they replicate in neuronal nuclei, establish long-term non-cytolytic infections, alter neural progenitor differentiation ^8^, and leave extensive genomic traces in the form of endogenous bornavirus-like elements (EBLs) ^9^. These viral “fossil records” provide a rare opportunity to reconstruct ancient host-virus interactions, and to map their evolution against encephalization trends.

The aim of this study is to investigate the potential of BDV as a driver of brain size evolution across a large number of mammalian species, within the paradigm of symbiotic virus-host coevolution ^10^. We first asked whether EBL loads across mammalian taxa predict observed encephalization trends, independently from pallium neuron count and maximum lifespan. We then devised a model to evaluate the potential importance of persistent neurotropic viral pressure in rapid developmental changes such as the hominin-specific fixation of *ARHGAP11B*, a gene that has recently been proven central to neocortex expansion ^11^. We eventually discuss whether this toy example can serve as a template for a wider range of brain evolution patterns through vertebrate history.

### BDV and encephalization trends

Borna disease virus (BDV) is a non-retroviral, highly neurotropic RNA virus, and persists in the central nervous system of infected individuals for their entire life span ^12^. Infected hosts develop a wide spectrum of neurological disorders, ranging from immune-mediated disease to behavioral alteration without inflammation, including deficits in learning and social behavior ^12;13^. It has been shown that BDV infects human neural progenitor cells (hNPC) and impairs neurogenesis ^14^. Unlike other typical neurotropic viruses, bornaviruses replicate noncytopathically and establish long-lasting, persistent infections in the nuclei of cultured cells and various tissues ^15^. Even relatively old EBLs might inhibit the replication of modern viruses ^9^, which indicates that species with a higher EBL count likely share a long common history with the virus. Although some EBLs are known to have been co-opted by their hosts ^16^, no direct link between EBL integrations and brain functions has been established so far.

Using data from the macroevolutionary history of bornaviral integration events of the last 100 Mya retraced by Kawasaki et al. ^17^, we tested the potential correlation between *encephalization slopes* (ES) as calculated by Shultz and Dunbar, and the mean number of EBL integrations in host genomes within previously studied taxa: Anthropoids, Strepsirrhines, Perissodactyls, Odontoceti, Artiodactyls, Feliformes, and Caniformes.

Building on recent analyses of natural and virtual endocasts ^18–23^, we extended the scope of Shultz and Dunbar’s encephalization slopes to include four supplementary taxa: the order Proboscidae, the suborder Sciuromorpha, the infraorder Hystricognathi and the family Hipposideridae. This allowed us to include taxa from two orders (Rodentia and Chiroptera) where evidence for the Social Brain Hypothesis is notably lacking ^24^. To avoid confounding effects arising from the considerable variation in rodent sociality, we focused exclusively on the obligately social representatives of both Sciuromorpha and Hystricognathi.

To account for other factors potentially influencing EBL integration events, we used AnAge and other sources from the literature to map correlations between EBL count and maximum longevity, and EBL count and total pallium neuron number.

#### Results

We found a strong correlation between encephalization slopes and mean EBL integration numbers per taxa (0.90; P ≤0.001) (**Table 1** and **Figure 1**).

**Table 1:** Encephalization slopes over time and EBL count.

| Taxon | E.Slope | P | Mean EBL nb |
| --- | --- | --- | --- |
| Anthropoids | 0.0160 | $\leq 0.001$ | 34 |
| Proboscidae | 0.0156 | 0.003 | 24 |
| Strepsirrhines | 0.0090 | 0.004 | 8 |
| Perissodactyls | 0.0086 | $\leq 0.001$ | 7 |
| Odontoceti | 0.0060 | $\leq 0.03$ | 4.57 |
| Hipposideridae | 0.0048 | 0.001 | 3 |
| Caniformes | 0.0043 | 0.003 | 4.2 |
| Artiodactyls | 0.0038 | 0.31 | 5 |
| Hystricognathi | 0.0018 | 0.003 | 4.8 |
| Feliformes | 0.0010 | 0.378 | 4 |
| Sciuromorpha | -0.0002 | 0.002 | 1 |

**Figure 1:**
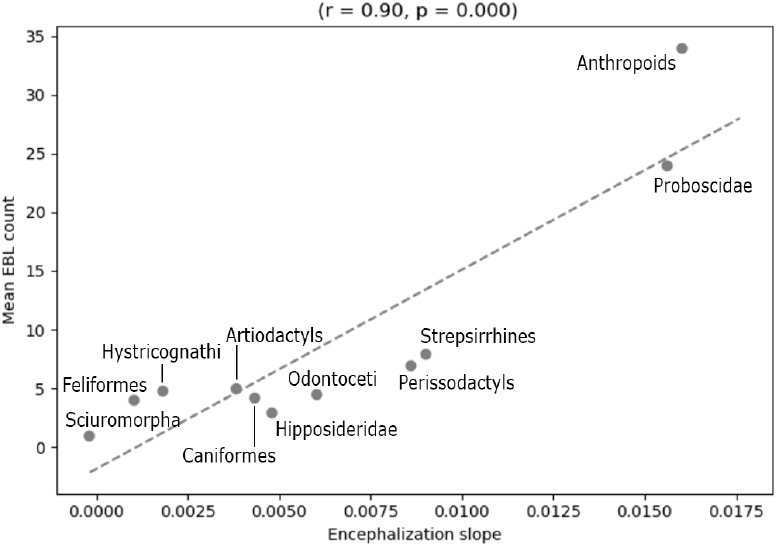
EBL integrations and encephalization slopes

To assess whether the addition of these four taxa affected the correlation between sociality and encephalization slopes, we calculated their sociality index (proportion of stable societies) and merged it with the existing data.

Adding Hipposideridae, Proboscidae, and social Sciuromorpha and Hystricognathi notably degraded the correlation between encephalization slopes and proportion of stable societies (from R = 0.767, P = 0.008 to R = 0.44, P = 0.175). **(Figure 2)**

**Figure 2:**
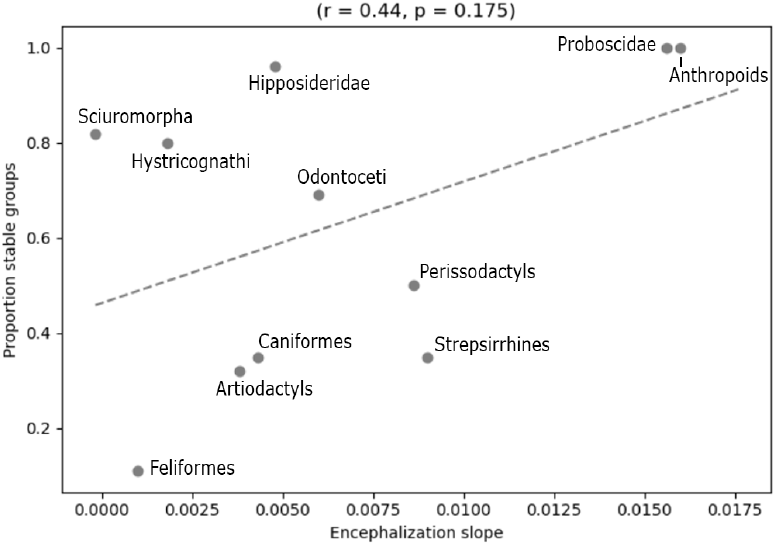
Proportion of stable groups and encephalization slopes

No meaningful correlation was found (r=0.335) between EBL integration numbers and maximum longevity. No correlation was found at the order level. A moderate correlation (r=0.477), at the macroscale level, was found between EBL integration numbers and total pallium neuron count. This correlation didn’t persist at the order level. As expected, EBL integrations don’t seem to necessitate shorter generations, large brains nor long lifespans. In fact, vastly different species such as *Rattus Norvegicus* or *Loxodonta Africana* have similar EBL counts.

The newly integrated data from fossils of these four taxa further suggests that sociality alone is not sufficient to drive encephalization; interestingly though, every lineage that underwent fast encephalization shows evidence of past bornaviral pressure, with social stability as a prerequisite.

However, the observed correlation does not prove causation, as there exist many other potential drivers of selective sweeps: environmental stress events, such as climate change ^25^, evolving patterns of predation ^26^, or uncontrolled demographic growth ^27^.

We thus proceeded to test different fixation scenarios on diploid Wright-Fisher simulations. As signatures of hard sweeps are most detectable when selection is relatively strong and recent (Kaplan, 1989) ^28^, we focused on the hominin-specific gene *ARHGAP11B*, whose emergence between 5 and 1 Mya drove a marked expansion of basal progenitors and neocortical growth. This gene provides a concrete test case for evaluating whether neurodevelopmental innovations could emerge primarily as compensatory mechanisms under neurotropic viral stress.

### Evaluating the influence of bornaviral pressure on the fixation time of a neurogenesis-enhancing gene

ARHGAP11B originated from a partial duplication of the ancestral *ARHGAP11A* gene and a subsequent splice site mutation that created a novel, human-specific C-terminal sequence ^29^. Functional experiments in mouse ^30^, ferret ^31^, and primate models ^32^ have shown that ARHGAP11B increases basal progenitor proliferation, expands the subventricular zone, and promotes neocortical folding, all traits that directly amplify neuron number during development. Despite its recent origin in the late Miocene or early Pliocene, *ARHGAP11B* is fixed in all modern humans, with no common gene-disruptive mutations in the general population ^33^.

Although the allele confers substantial neurogenic effects experimentally ^34^, cognitive and behavioral traits are complex, and selection on such traits in hominins is generally expected to be weak (*s* ≈ 10^−3^), diffusely expressed, thus producing long fixation times and frequent early loss ^35–37^. Under such weak selection, a beneficial allele arising once in a population of size *N*_*e*_ ≈ 1–5 × 10^4^ is predicted to be lost ∼ 99.8% of the time (*P*_fix_ ≈ 2*s*), and to fix over 200 kya when it does succeed; far slower than the empirical pattern. This discrepancy motivates the exploration of selective regimes capable of producing fast, hard sweeps under realistic demographic conditions.

In a hypothetical scenario in which most early hominins lacked ARHGAP11B and were chronically or episodically infected with BDV, ARHGAP11B expression could act as a partial compensatory mechanism by increasing the production of neural progenitors and extending neurogenesis. This might offset some neuron loss caused by BDV-induced apoptosis of newly formed neurons ^14^. However, ARHGAP11B cannot prevent BDV from killing differentiating neurons, so it cannot provide a complete rescue.

We implemented diploid Wright-Fisher simulations using parameters informed by ancestral hominin demography, with one scenario exploring the controversial ^38^ 1-Mya bottleneck hypothesis by Hu et al ^39^. Although our model simplifies demographic structure, age distribution, and migration, its conclusions concern the relative effects of selective regimes rather than precise sweep durations.

We modeled the evolution of a newly arisen beneficial allele under selective regimes differing in strength, temporal structure, and epidemiological context. Weak cognitive or behavioral advantages were represented by a constant selection coefficient *s*_cog_.

Bornavirus-like developmental pressure was modeled as episodic selection, in which infected cohorts experienced a fitness cost *c* and viral prevalence *q* fluctuated (see *Methods*). The model allowed the carriers of one or two alleles to mitigate this fitness cost by a factor *m*_*AA*_ or *m*_*Aa*_ = 0.8 ∗ *m*_*AA*_ (quasi-dominance), respectively. Genetic drift was taken into account.

We evaluated both single-origin mutations (*p*_0_ = 1*/*(2*N*_*e*_)) and alleles present as low-frequency standing variation, and for each condition we calculated fixation probabilities, mean fixation times, and mean loss times across 1,000 replicate simulations over a grid of variables spanning biologically plausible ranges.

To visualize how bornavirus-like pressure alters the trajectory of a neurogenesis-enhancing allele, we plotted multiple selective regimes on the same set of axes, allowing their dynamics to be compared directly (**Figures 3A-F**). Each panel displays allele-frequency trajectories for four conditions: (i) cognitive selection alone acting on single-origin mutations, (ii) cognitive selection acting on alleles present as low-frequency standing variation *p*_0_, (iii) bornavirus-like episodic selection acting on single-origin mutations, and (iv) bornavirus-like selection acting on alleles present as low-frequency standing variation. Each trajectory represents a single replicate drawn from a larger ensemble.

**Figure 3:**
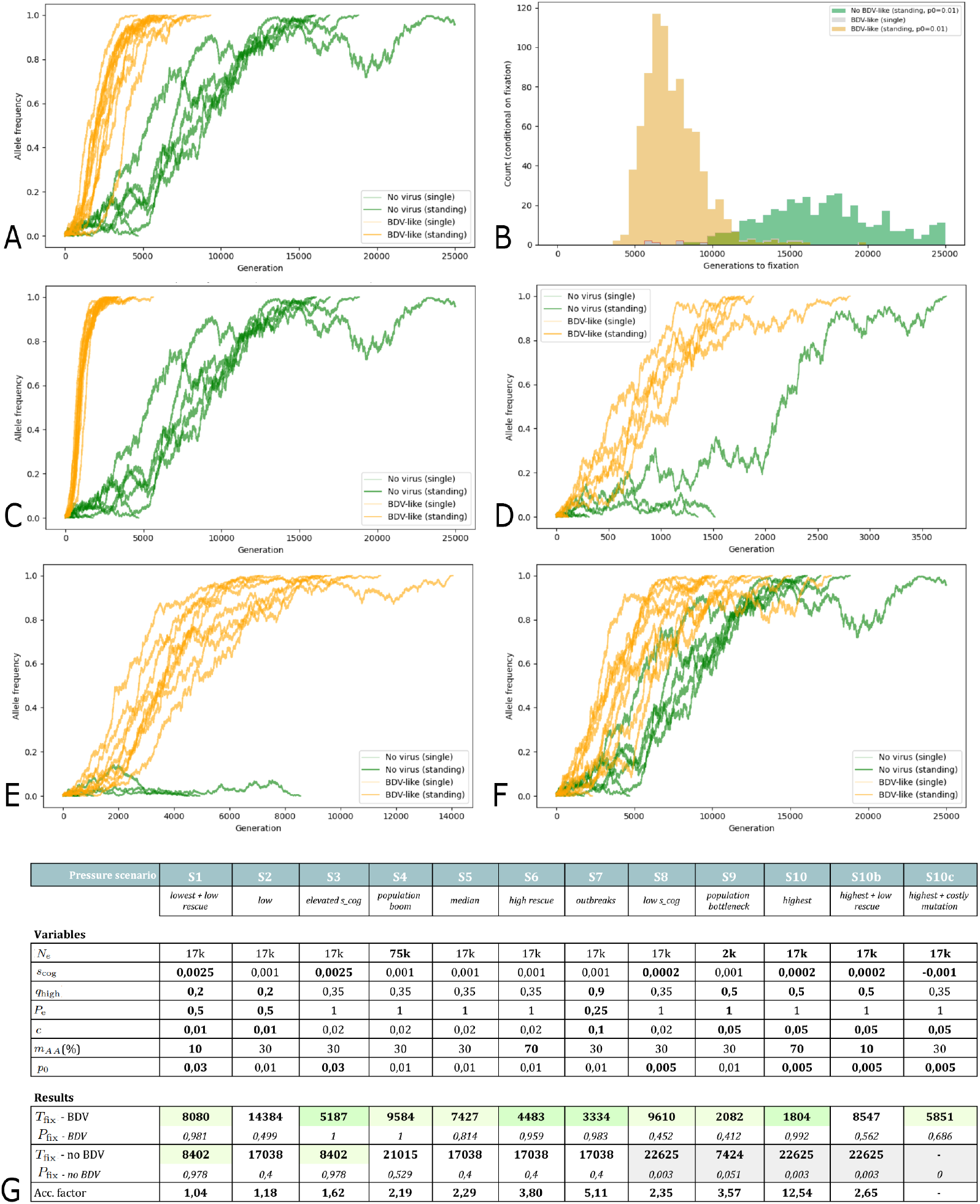
Evolutionary dynamics of an ARHGAP11B-like allele under BDV-like selective pressures. **(A)** Allele-frequency trajectories for **Scenario 5** (median parameters), showing faster fixation, and a doubled probability of fixation, under BDV-like pressure (orange) compared with no-virus conditions (green). **(B)** Distribution of fixation times for **Scenario 5**, contrasting BDV-like (orange) and no-virus (green) environments. **(C)** Allele-frequency trajectories for **Scenario 7**, illustrating the influence of intermittent, strong outbreaks on fixation times. **(D)** Allele-frequency trajectories for **Scenario 9** showing how loss due to a population bottleneck can be averted through bornaviral pressure. **(E)** Allele-frequency trajectories for **Scenario 8**, depicting conditions where the allele offers very low cognitive advantage. **(F)** Allele-frequency trajectories for **Scenario 2** depicting low bornaviral pressure. **(G)** Summary table of parameter sets (S1–S10c) and outcomes. In bold font: tweaked variables. Scenarios 1 and 10 represent, respectively, the least and the most favorable parameter settings for the BDV-pressure hypothesis. Light green shading indicates *possible* evolutionary scenarios; green shading indicates *plausible* scenarios consistent with biological constraints. Light grey cells indicate quasi-certain allele loss. *T*_fix_: mean fixation time, in generations; *P*_fix_: fixation probability. *Acc. factor* : acceleration factor under BDV-like pressure. For the other variables, see main text.

Twelve from the most significant scenarios are shown on the corresponding table (**Figure 3G**), of which five are plotted.

#### Results

In our simulations, neurotropic bornaviral-like pressure accelerates the fixation of the neurogenesis-promoting gene; in some cases, it allows it. In most scenarios, the fixation probability in notably increased under BDV-like pressure.

Fixation was very unlikely (*P*_fix_ *<* 0.005) for single-origin mutations in all scenarios. Standing variation, indicating a long-term, weak advantage, was necessary in all cases. This advantage could stem from pre-existing cognitive benefits, prior viral pressure, or a combination thereof.

The scenario with median values **(S5)** reduces fixation time by roughly two-fold, from over 17,000 generations (425 kya) to under 7,500 generations (180 kya), and displays a two-fold heightened fixation probability.

When the virus has low prevalence and exerts low fitness cost **(S1-S2)**, or when *s*_cog_ is assumed to be high **(S3)**, the bornaviral pressure hypothesis does not offer any meaningful advantage.

However, when *s*_cog_ is low or negative **(S8, S10, S10b, S10c)**, bornaviral pressure is the only path to fixation.

Generally, a high BDV prevalence and a marked fitness cost strongly promote allele fixation. Scenario 10b shows that under certain circumstances, even a very moderate compensatory mechanism (10% rescue only) makes fixation possible.

While scenarios 10a, b and c may look extreme when looking at current BDV exposure and effects in contemporary humans, their 5% cost and seasonal pattern is based on what is observed for PaBV (Parrot Bornavirus) in some species of caged parrots ^40^. As a side note, although parrots are not mammals, they have remarkable brains for their size ^41^, and many suffer from stereotypy (abnormal repetitive behavior) that some studies have linked to an abnormal developmental process, similar to what is observed in human autistic or untreated schizophrenic patients ^42^. Since an association between BDV and schizophrenia is suspected ^43^, further research on the links between avian bornaviruses and neurodevelopmental burden could be of interest. It could be that caged parrots are experiencing a sustained neurotropic bornaviral pressure similar to what has triggered the integration of multiple EBLs in primate lineages – unfortunately, the absence of LINE-1 machinery ^44^ (which seems responsible for the endogenization of bornaviral elements ^8^) in birds makes their co-evolution history with bornaviruses arduous to study.

On the other end of the spectrum, rare and lethal outbreaks of Borna Disease are sometimes observed in horses or sheep. Mixing high seroprevalence in endemic regions (11.5-22.5%), very low disease incidence (0.02-0.04%), but high mortality (90%) ^45^, this regime is unlikely to have a genomic impact. Scenario 7 illustrates an intermediate case between PaBV-like seasonal infections and such mammalian bornaviral outbreaks. Our simulation shows that a 10% cost on fitness every four generations can still lead to *ARHGAP11B*-like fixation.

## Discussion

Our analyses of the potential role of the neurotropic Borna Disease Virus (BDV) in mammalian encephalization point to virus-mediated stress as a factor that may merit further consideration within established evolutionary paradigms.

First, recent data from fossil species allowed us to expand Shultz and Dunbar’s analysis of encephalization slopes to four new orders, and notice a correlation between the amount of endogenous bornaviral-like elements (EBLs) and encephalization rate, with stable sociality as a prerequisite.

Secondly, our simulations of different fixation scenarios involving *ARHGAP11B* as a compensatory mechanism under realistic, biologically informed bornaviral pressure painted a picture of a complex evolutionary landscape where viral-host coexistence might be mutually beneficial in the long run. Although the dichotomy between viral and non-viral environmental stress might appear too straightforward in these simulations, in practice these pressures were likely to have fluctuated and overlapped through time; for instance, it is plausible that bornaviral pressure helped buffer the additional metabolic and obstetric costs during the early phase of allele spread, and that later, subsequent organismal changes made the allele beneficial even without viral stress.

These interpretations must be considered alongside several limitations. Firstly, despite recent advances in EVE detection, and growing interest in non-human paleoneurobiology, data is still scarce. Encephalization trends must be taken with extra caution; new techniques and measurements will likely add more species to our corpus in the future.

Although we looked at brain size and longevity, other confounding factors such as genome assembly quality, retrotransposon activity, and effective population size might influence the observed EBL count. In particular, it is not very clear why separate taxa within the orders Rodentia and Chiroptera have very different EBL counts; this may reflect lineage-specific differences in immune strategies.

Regarding cognition, the brain of birds and bats might need specific metrics, like neuron density, connectivity, or renewal, to take the strong ecological constraints limiting their weight into account.

Our simulations extrapolate from current bornaviral events, and to date, there is no way to know the exact etiology and pathophysiology of ancient mammalian bornaviruses, although it might be possible soon ^46;47^. Nevertheless, given the non-cytolytic, sublethal nature of bornaviruses in species that tolerate them, it would already be informative to investigate their effect on supposedly healthy subjects ^48;49^.

Lastly, we focused on viral pressure alone, and did not look at EBLs as promoters of molecular mechanisms having an influence on cognition or other traits.

Yet, taken together, the insights from aforementioned correlations and simulations allow us to propose the *Neurotropic Viral Pressure Hypothesis* (**Figure 4**): where bornaviral-like infections act as accelerators and mediators within the classical Socio-ecological brain-encephalization loop.

**Figure 4:**
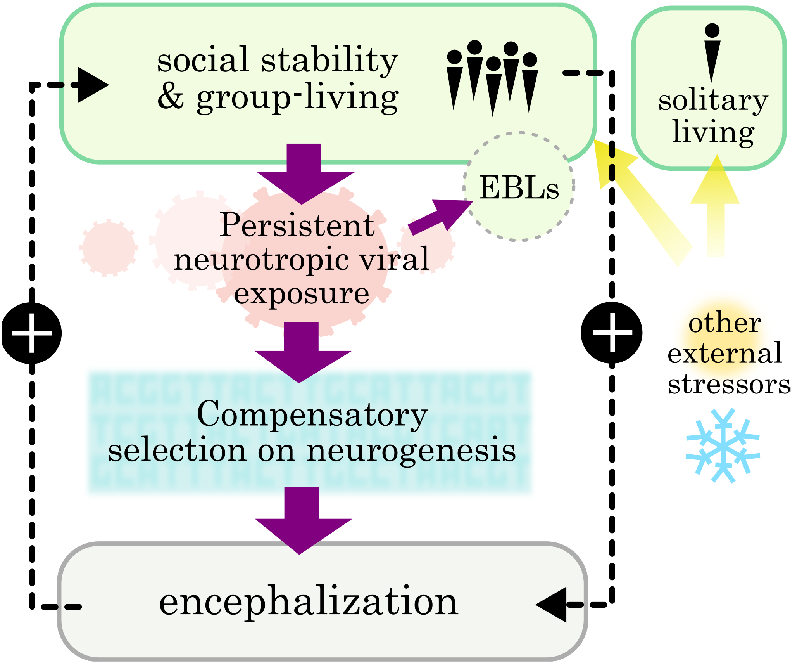
The Neurotropic Viral Pressure Hypothesis.

It is important to note that this hypothesis is not at odds with ecological theories of encephalization. Climate changes, ecological disasters and massive extinctions usually go hand in hand with increased virus circulation: bats, as an example, are particularly sensitive to climate change due to their high surface-to-volume ratio and low reproductive rates ^50^; their migrations are linked to interspecies epidemic events. However, environmental stress alone does not target the brain specifically. While food scarcity impedes neurogenesis as well ^51^, it impacts every organ of the body, thus accelerating the positive selection of multiple features, homothetically; whereas BDV pressure likely favors brain-oriented mutations. And if EBLs reflected genomic turbulence alone, they would be randomly distributed across clades.

It is known that in times of rapid environmental change, social groups have the ability to heavily influence selection by outcasting less able individuals ^52^. That behavior could include the negative selection of detrimental cognitive traits; but again, this is fully compatible with the NVP hypothesis, since this kind of active social regulation increases the cost associated with BDV-induced brain impairments.

Sustained neurotropic viral pressure could thus explain why encephalization occurred only in social taxa during such ecological stress events, and how similar species could have diverged dramatically in terms of encephalization; and conversely, why very different groups converged across continents during short saltational events.

A practical way to evaluate the ideas developed here would be to test whether ARHGAP11B (over-)expression can partially rescue BDV-induced neural phenotypes in a controlled experimental system.

The case of parrots, and birds more broadly, merits further exploration. Although this study focuses on mammals, there is no reason to think that the principles described here could not be extended to other classes.

More broadly, virus-host coevolution is often interpreted through the lens of immunity and framed as an escalatory “arms race”. Yet many other modalities of interaction are possible, including cases in which persistent viral pressure favors developmental innovation rather than enhanced immunity per se. Considering these alternative pathways of virus-mediated selection may therefore broaden our understanding of how environmental, developmental, and infectious pressures intersect to influence organismal evolution across vertebrates.

## Methods

### 1. Data Cleaning

The AnAge longevity dataset and the endogenous bornavirus-like element (EBL) count dataset from Kawasaki et al. (2021) ^17^ were cleaned and standardized prior to analysis. Raw AnAge CSV files contained irregular quoting, so each line was trimmed, leading quotation marks were removed, and trailing quote–semicolon artifacts were stripped. The corrected file was then parsed to extract taxonomic and biological fields, and numeric columns (adult body mass and maximum longevity) were coerced into numeric format. Scientific names were reconstructed by concatenating genus and species fields. For the EBL dataset, the supplemental XLSX file was imported and the true header row was identified programmatically by locating the “Host species” label. Only the EBL name and host species fields were retained. Viral genus was extracted from the EBL identifier using regular-expression parsing, and counts of EBLs were aggregated by host species and viral genus before being pivoted into a wide table. To harmonize taxonomic identifiers across datasets, host species names and AnAge scientific names were normalized by lowercasing, removing punctuation, replacing underscores, and trimming whitespace. The cleaned datasets were then used for downstream comparative analyses, which were all performed in Python.

### 2. Estimating Encephalization Rates for Newly Added Taxa

For Sciurmorpha, Hystricognathi, Proboscidea, and Hipposideridae, we assembled fossil and extant estimates of brain mass, body mass, and geological age (Mya) into taxon-specific datasets, from the following sources: Sciuromorpha ^19;20^, Hystricognathi ^19;20;22;23^, Proboscidae ^18^ and Hipposideridae ^21^. When needed, endocranial volumes of rodents were converted to mass by dividing the volume by 1.05^54^. For extant species, we curated the Dryad dataset and created four separate CSV files. Following the approach of Shultz and Dunbar (2010) ^5^, we quantified each taxon’s encephalization rate as the temporal trend in relative brain size.

Brain and body masses were log-transformed, and we fitted the multiple regression model

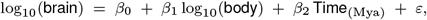

where *β*_0_ is the intercept and *ε* is the residual error term. The coefficient associated with time, *β*_2_, was taken as the *encephalization slope* for the taxon, and its associated *P*-value was used to assess whether the temporal trend differed significantly from zero.
Shultz and Dunbar (2010) estimated temporal slopes using phylogenetic generalized least squares (PGLS), but also reported slopes from nonphylogenetic general linear models (GLMs). These two approaches were found to be highly correlated (order level: *R* = 0.83, *P* = 0.04; suborder level: *R* = 0.89, *P* = 0.001), indicating that both yield comparable encephalization-rate estimates. Because fossil datasets for the newly added taxa lack resolved phylogenies of fossil species, we followed the GLM implementation.

Because Proboscidae includes only two extant species compared with eight fossil species, we generated additional synthetic datapoints based on the observed encephalization quotients to balance representation within the clade.

We focused on social species within the suborder Sciuromorpha (mostly ground squirrels and marmots), and the infraorder Hystricognathi (naked mole rats, chinchillas, Caviidae). See *Data and Code Availability* for the detail.

### 3. Comparative Analyses of Encephalization Rate, EBL Burden, and Sociality

We assembled a taxon-level dataset combining encephalization slopes (estimated as described above), mean endogenous bornavirus-like element (EBL) counts per taxon (using the data extracted at step 1), and the proportion of extant species living in stable social groups. Because EBL burdens vary only within a moderate numerical range across clades (1–45 insertions) and do not exhibit strong overdispersion, we conducted analyses on raw counts rather than log-transformed values.

The taxa Hipposideridae and Proboscidea were represented by a single species with documented EBL counts, but this is unlikely to bias clade-level means because EBL burden shows low within-clade variation across the broader dataset, with closely related species typically differing by only a few percentage points.

We characterized each extant species of the four new taxa according to two social categories: (i) solitary + facultatively social, and (ii) stable social groups. Species typical behavior was taken from the Animal Diversity Web (https://animaldiversity.org). For each taxon, the proportion of species that demonstrated stable social groups was calculated.

For each taxon, we tested the association between encephalization slope and (i) mean EBL count and (ii) the proportion of stable societies using Pearson correlation, following the comparative framework described in step 2. Scatterplots with linear fits were generated for visualization. The data is stored in the notebook directly.

### 4. EBLs and other traits

We then asked whether EBL burden was really a hint of past bornaviral exposure, or a mere byproduct of some other traits. We first assessed whether maximum longevity alone could explain variation in EBL burden. Maximum lifespan (log-transformed) showed only a weak association with total EBL count (Pearson r = 0.335, p = 0.0004). Thus, lifespan is not a sufficient driver of EBL accumulation and cannot account for the much stronger correlation observed between EBL burden and encephalization rate.

The notebook *“EBL longevity control*.*ipynb”*, as a sidenote and for further exploration, contains the correlation between EBL count and longevity residual (due to the fact that we first started this study looking for an influence of endogenous bornavirus-like elements on longevity). Maximum longevity and adult body mass were *log*_10_-transformed, and mass-corrected longevity was computed as the residuals of an ordinary least squares regression of log longevity on log mass. These residuals represent lifespan relative to that expected for a mammal of the same body size. Residual values were merged with EBL counts using the data from step 1. For each mammalian order containing at least four species, we tested for associations between longevity residuals and total or genus-specific EBL counts using Spearman rank correlations. No correlation was found.

Also, to test whether EBL numbers could be explained by present-day brain size, we conducted an independent analysis using published pallium neuron counts ^55^. Pallium neuron numbers were *log*_10_-transformed to account for the several-order-of-magnitude variation across mammals and merged with species-level EBL counts. Associations between log neuron count and total or genus-specific EBL numbers were assessed using Spearman rank correlations. This yielded a moderate correlation (rho=0.477, p=0.00246). Labelling each species extensively allowed us to spot taxa and notice no correlation within them. While EBLs are obviously more frequent in big-brained species, the correlation with the encephalization slope is much stronger.

As a sidenote again, Bornaviruses include several genera (Orthobornavirus, Carbovirus, Cultervirus). We have collapsed them for clarity, but Carboviruses show an intriguing correlation with neuron count.

### 5. Fixation scenarios

We modelled the evolutionary dynamics of a beneficial allele in a hominin-like population using a diploid Wright– Fisher model with viability selection and genetic drift. Each simulation tracked allele-frequency change in a population of effective size *N*_*e*_ with discrete, non-overlapping generations. Genotype fitness depended on two components: (i) a baseline cognitive advantage *s*_cog_ conferred by the derived allele, and (ii) a virus-related fitness penalty proportional to the environmental viral prevalence *q*_*t*_.

In each generation, viral prevalence *q*_*t*_ was, in principle, drawn from a two-state model (baseline *q*_low_ vs. epidemic *q*_high_) with probability *P*_e_, via *q*_*t*_ ∈ { *q*_low_, *q*_high_ }. The ancestral genotype suffered a fitness penalty *q*_*t*_ ∗ *c*, whereas heterozygotes and homozygotes for the derived allele experienced reduced penalties (through mitigation factors for viral damage) and a cognitive fitness advantage of 0.8 *s*_cog_ and *s*_cog_, respectively.

After computing genotype-specific fitnesses, we calculated the post-selection allele frequency and obtained the next generation’s allele count by binomial sampling of 2*N*_*e*_ gametes, following the standard Wright-Fisher sampling process. We simulated two introduction regimes for the derived allele: (i) a single new mutation (*p*_0_ = 1*/*(2*N*_*e*_)), and (ii) standing variation *p*_0_. For each regime, we ran scenarios with and without BDV-like viral pressure (no-virus scenarios used *q*_low_ = *q*_high_ = 0 and *c* = 0). For each scenario, we ran 1,000 replicate trajectories for up to 25,000 generations and recorded the probability of fixation and the distribution of fixation times (conditional on fixation).

*N*_*e*_ : we chose 17k as the main value ^53^, but tested scenarios with 2k (population bottleneck) and 75k (population boom).

*s*_cog_ : We chose 0.001 as the median value, consistent with empirical estimates for complex behavioral traits, but we also tested scenarios where the selection for cognition was stronger, and others where it was negligible and even negative, reflecting increased obstetric or metabolic costs. *p*_0_ (standing genetic variation) was correlated to *s*_cog_, and its value ranged from 0.5% to 3%, with 1% as the main estimation.

*c* was estimated to vary between 0.01 (1% reduction in fitness) and 0.1 in the case of lethal, centennial outbreaks; however, values of 0.02 and 0.05 were often preferred. These estimates are based on data from the literature ^56–58^.

*P*_e_, *q*_low_ and *q*_high_ vary according to different regimes corresponding to real-world data about mammalian and avian bornaviruses ^56–58^. We chose to focus on bornaviruses not only because EBLs are well documented, but also because BDV has specific modes of infection that make it rather “silent”. It could be argued that numerous neurotropic viruses might exert similar evolutionary pressures on the brain. Viruses such as rabies, Nipah, Hendra, West Nile or HSV-1 may be too lethal, too sporadic, or too transient in their effects to plausibly act as long-term selective forces; however, for future studies, it could be interesting to look at viruses that typically target one organ and to see whether regularly infected species have developed genetic countermeasures. *m*_*AA*_ was estimated at around 30% (with scenarios at 10% and 70%), consistent with in vivo and in vitro experiments ^14;30^. A full rescue of the viral cost *c* was considered unlikely.

In our model, we have not considered the observation that the BDV X protein can protect neurons from degeneration in tissue culture and in animal models of Parkinson’s disease and ALS ^48;59^, as part of the virus’s strategy to maintain replication in neurons without causing cell death. Such late-life neuroprotective effects are unlikely to influence host evolution, because selection on traits expressed after reproduction is minimal. Nevertheless, they remain noteworthy from a viral perspective: extending neuronal survival in older hosts increases the duration of infection and thus the reservoir available for viral maintenance and transmission, even when individuals are no longer reproductively active.

## Data and code availability

All data are available in the main text or at https://github.com/EmmanuelGauvain/NVPh_2025. Custom code used in this study is available at the same URL.

## Acknowledgements

The author would like to thank Amont Biosciences, M.v.E., B.A.y A., A.F., M.d.V., D.M. and M.R.

## References

[1] Kaufman JA (2003) On the expensive tissue hypothesis: independent support from highly encephalised fish. Curr Anthropol 44:705–706.

[2] Allman J, McLaughlin T, Hakeem A (1993) Brain weight and lifespan in primate species. Proc Natl Acad Sci USA 90:118–122. doi:10.1073/pnas.90.1.118

[3] Hardie JL, Cooney CR (2023) Sociality, ecology and developmental constraints predict variation in brain size across birds. J Evol Biol 36:144–155. doi:10.1111/jeb.14117

[4] Byrne RW, Whiten A (1988) Machiavellian intelligence: social expertise and the evolution of intellect in monkeys, apes and humans. Rev Philos France É tranger 179:627–628.

[5] Shultz S, Dunbar R (2010) Encephalization is not a universal macroevolutionary phenomenon in mammals but is associated with sociality. Proc Natl Acad Sci USA 107:21582–21586. doi:10.1073/pnas.1005246107

[6] Smaers JB, Dechmann DKN, Goswami A, Soligo C, Safi K (2012) Comparative analyses of evolutionary rates reveal different pathways to encephalization in bats, carnivorans and primates. Proc Natl Acad Sci USA 109:18006–18011. doi:10.1073/pnas.1212181109

[7] Shultz S, Nelson E, Dunbar RIM (2012) Hominin cognitive evolution: identifying patterns and processes in the fossil and archaeological record. Philos Trans R Soc Lond B Biol Sci 367:2130–2140. doi:10.1098/rstb.2012.0115

[8] Honda T (2022) Bornavirus infection in human diseases and its molecular neuropathology. Clin Exp Neuroimmunol 13:7–16. doi:10.1111/cen3.12686

[9] Horie M, Kobayashi Y, Suzuki Y, Tomonaga K (2013) Comprehensive analysis of endogenous bornavirus-like elements in eukaryote genomes. Philos Trans R Soc Lond B Biol Sci 368:20120499. doi:10.1098/rstb.2012.0499

[10] Roossinck MJ, Bazán ER (2017) Symbiosis: viruses as intimate partners. Annu Rev Virol 4:123–139. doi:10.1146/annurev-virology-110615-042323

[11] Florio M et al. (2016) A single splice site mutation in human-specific ARHGAP11B causes basal progenitor amplification. Sci Adv 2:e1601941. doi:10.1126/sciadv.1601941

[12] Solbrig MV, Koob GF (2003) Neuropharmacological sequelae of persistent CNS viral infections: lessons from Borna disease virus. Pharmacol Biochem Behav 74:777–787. doi:10.1016/S0091-3057(03)00019-4

[13] Hornig M et al. (2001) Borna disease virus infection of adult and neonatal rats: models for neuropsychiatric disease. Curr Top Microbiol Immunol 253:157–177.

[14] Brnic D et al. (2012) Borna disease virus infects human neural progenitor cells and impairs neurogenesis. J Virol 86:2512–2522. doi:10.1128/JVI.05663-11

[15] Schneider U, Martin A, Schwemmle M, Staeheli P (2007) Genome trimming by Borna disease viruses. Cell Mol Life Sci 64:1038–1042.

[16] Kobayashi Y et al. (2016) Exaptation of bornavirus-like nucleoprotein elements in Afrotherians. PLoS Pathog 12:e1005785. doi:10.1371/journal.ppat.1005785

[17] Kawasaki J et al. (2021) A 100-My history of bornavirus infections hidden in vertebrate genomes. Proc Natl Acad Sci USA 118:e2026235118. doi:10.1073/pnas.2026235118

[18] Benoit J et al. (2019) Brain evolution in Proboscidea across the Cenozoic. Sci Rep 9:9323. doi:10.1038/s41598-019-45888-4

[19] Bertrand OC, Amador-Mughal F, Silcox MT (2016) Virtual endocasts of Eocene Paramys (Paramyinae): oldest endocranial record for Rodentia and early brain evolution in Euarchontoglires. Proc Biol Sci. 2016 Jan 27;283(1823):20152316. doi:10.1098/rspb.2015.2316

[20] Bertrand OC, Silcox MT (2023) Brain evolution in fossil rodents: a starting point. In: Dozo MT et al. (eds) Paleoneurology of Amniotes. Springer, Cham. doi:10.1007/978-3-031-13983-316

[21] Yao L et al. (2012) Evolutionary change in the brain size of bats. Brain Behav Evol 80:15–25. doi:10.1159/000338324

[22] Ferreira JD, Rinderknecht A, de Moura Bubadué J, Gasparetto LF, Dozo MT, Sánchez-Villagra MR, Kerber L. (2024) Unveiling the neuroanatomy of Josephoartigasia monesi and the evolution of encephalization in caviomorph rodents. Brain Struct Funct. 2024 May;229(4):971–985. doi:10.1007/s00429-024-02762-y

[23] Fernández Villoldo JA, Verzi DH, Olivares AI, Dos Reis SF, Lopes RT, Perez SI. (2025) Exploring the palaeoneurology of the extinct spiny rat *Eumysops chapalmalensis* (Hystricognathi: Echimyidae): a comparative phylogenetic analysis of brain size and shape. Zoological Journal of the Linnean Society. 203(3): zlaf005. doi:10.1093/zoolinnean/zlaf005

[24] Kverková K et al. (2018) Sociality does not drive the evolution of large brains in eusocial African mole-rats. Sci Rep 8:9203. doi:10.1038/s41598-018-26062-8

[25] Bonelli S et al. (2011) Population extinctions in Italian diurnal Lepidoptera. J Insect Conserv 15:879–890. doi:10.1007/s10841-011-9387-6

[26] Nair RR et al. (2019) Bacterial predator–prey coevolution accelerates genome evolution. Nat Commun 10:4301. doi:10.1038/s41467-019-12140-6

[27] Wilson BA, Petrov DA, Messer PW (2014) Soft selective sweeps in complex demographic scenarios. Genetics 198:669–684. doi:10.1534/genetics.114.165571

[28] Kaplan NL, Hudson RR, Langley CH (1989) The hitchhiking effect revisited. Genetics 123:887–899.

[29] Florio M et al. (2015) Human-specific ARHGAP11B promotes neocortex expansion. Science 347:1465–1470. doi:10.1126/science.aaa1975

[30] Xing L et al. (2021) Expression of ARHGAP11B in mice leads to neocortex expansion. EMBO J 40:e107093. doi:10.15252/embj.2020107093

[31] Kalebic N et al. (2018) ARHGAP11B induces neocortical expansion in ferrets. eLife 7:e41241. doi:10.7554/eLife.41241

[32] Heide M et al. (2020) ARHGAP11B increases size and folding of primate neocortex. Science 369:546–550. doi:10.1126/science.abb2401

[33] Dennis MY et al. (2017) Human-specific segmental duplications. Nat Ecol Evol 1:0069. doi:10.1038/s41559-016-0069

[34] Fischer J et al. (2022) ARHGAP11B ensures humanlike progenitor levels in organoids. EMBO Rep 23:e54728. doi:10.15252/embr.202254728

[35] Sella G, Barton NH (2019) Thinking about the evolution of complex traits. Annu Rev Genomics Hum Genet 20:461–493. doi:10.1146/annurev-genom-083115-022316

[36] Zeng J et al. (2021) Widespread signatures of natural selection. Nat Commun 12:1164. doi:10.1038/s41467-021-21446-3

[37] Kimura M (1962) On the probability of fixation of mutant genes. Genetics 47:713–719.

[38] Cousins T, Durvasula A (2025) Insufficient evidence for a severe human bottleneck. Mol Biol Evol 42:msaf041. doi:10.1093/molbev/msaf041

[39] Hu W et al. (2023) Genomic inference of a severe human bottleneck. Science 381:979–984. doi:10.1126/science.abq7487

[40] Villanueva BHA et al. (2024) Surveillance of parrot bornavirus in Taiwan. Viruses 16:805. doi:10.3390/v16050805

[41] Olkowicz S et al. (2016) Birds have primate-like numbers of neurons. Proc Natl Acad Sci USA 113:7255–7260. doi:10.1073/pnas.1517131113

[42] Garner JP, Meehan CL, Mench JA (2003) Stereotypies in parrots and links to psychiatric disease. Behav Brain Res 145:125–134. doi:10.1016/S0166-4328(03)00115-3

[43] Azami M et al. (2018) Association between Borna disease virus and schizophrenia. Asian J Psychiatr 34:67–73. doi:10.1016/j.ajp.2017.11.026

[44] Kapusta A, Suh A (2017) Evolution of bird genomes: a transposon’s-eye view. Ann N Y Acad Sci 1389:164–185. doi:10.1111/nyas.13295

[45] Piotrowski I, Hilbe M, Junge H (2025) Borna disease virus infection in horses and donkeys. Equine Vet Educ. doi:10.1111/eve.14220

[46] Callaway HM et al. (2019) Ancient endogenous parvovirus remnants in rodents. J Virol 93:e01542–18. doi:10.1128/JVI.01542-18

[47] Gifford RJ (2021) Mapping the evolution of bornaviruses. Proc Natl Acad Sci USA 118:e2108123118. doi:10.1073/pnas.2108123118

[48] Tournezy J, Léger C, Klonjkowski B, Gonzalez-Dunia D, Szelechowski M, Garenne A, Mathis S, Chevallier S, Le Masson G (2024). The Neuroprotective Effect of the X Protein of Orthobornavirus Bornaense Type 1 in Amyotrophic Lateral Sclerosis. IJMS, 25(23), 12789. doi:10.3390/ijms252312789

[49] Marty FH, Bettamin L, Thouard A, Bourgade K, Allart S, Larrieu G, Malnou CE, Gonzalez-Dunia D, Suberbielle E. (2021) Borna disease virus docks on neuronal DNA double-strand breaks to replicate and dampens neuronal activity. iScience. 2021 Dec 16;25(1):103621. doi:10.1016/j.isci.2021.103621

[50] Festa F et al. (2022) Bat responses to climate change. Biol Rev 98:1–21. doi:10.1111/brv.12893

[51] Galler JR et al. (2021) Neurodevelopmental effects of childhood malnutrition. NeuroImage 231:117828. doi:10.1016/j.neuroimage.2021.117828

[52] Foley C, Pettorelli N, Foley L (2008) Severe drought and calf survival in elephants. Biol Lett 4:541–544. doi:10.1098/rsbl.2008.0370

[53] Huff CD, Xing J, Rogers AR, Witherspoon D, Jorde LB (2010) Mobile elements reveal small population size in the ancient ancestors of Homo sapiens. Proc Natl Acad Sci USA. 107(5):2147–52. doi: 10.1073/pnas.0909000107. doi: 10.1073/pnas.0909000107

[54] Hofman MA. (1983) Encephalization in hominids: evidence for the model of punctuationalism. Brain. Behav. Evol. 122, 102–117. doi:10.1159/000121511

[55] Herculano-Houzel S, Catania K, Manger P Kaas J (2015). Mammalian Brains Are Made of These: A Dataset of the Numbers and Densities of Neuronal and Nonneuronal Cells in the Brain of Glires, Primates, Scandentia, Eulipotyphlans, Afrotherians and Artiodactyls, and Their Relationship with Body Mass. Brain, behavior and evolution. doi: 10.1159/000437413

[56] Bautista JR et al. (1994) Abnormalities in rats with neonatally acquired Borna virus infection. Brain Res Bull 34:31–40. doi:10.1016/0361-9230(94)90183-x

[57] Gonzalez-Dunia D, Sauder C, de la Torre JC (1997) Borna disease virus and the brain. Brain Res Bull 44:647–664. doi:10.1016/S0361-9230(97)00276-1

[58] Rubbenstroth D (2022) Avian bornavirus research: a comprehensive review. Viruses 14:1513. doi:10.3390/v14071513

[59] Szelechowski M, Bétourné A, Monnet Y et al (2014) A viral peptide that targets mitochondria protects against neuronal degeneration in models of Parkinson’s disease. Nat Commun 5:5181 doi:10.1038/ncomms6181

